# Machine learning-based investigation of the cancer protein secretory pathway

**DOI:** 10.1101/2020.09.09.289413

**Authors:** Rasool Saghaleyni, Azam Sheikh Muhammad, Pramod Bangalore, Jens Nielsen, Jonathan L. Robinson

**Affiliations:** Department of Biology and Biological Engineering, Chalmers University of Technology, SE-412 96 Gothenburg, Sweden; Department of Computer Science and Engineering, Chalmers University of Technology, SE-412 96 Gothenburg, Sweden; Greenbyte AB, Ostra Hamngatan 16, SE-41109 Gothenburg, Sweden; Wallenberg Center for Protein Research, Chalmers University of Technology, Kemivägen 10, SE-41258 Gothenburg, Sweden; BioInnovation Institute, Ole Maaløes Vej 3, DK-2200 Copenhagen, Denmark; Department of Biology and Biological Engineering, National Bioinformatics Infrastructure Sweden, Science for Life Laboratory, Chalmers University of Technology, Kemivägen 10, SE-41258 Gothenburg, Sweden.

## Abstract

Deregulation of the protein secretory pathway (PSP) is linked to many hallmarks of cancer, such as promoting tissue invasion and modulating cell-cell signaling. The collection of secreted proteins processed by the PSP, known as the secretome, is often studied due to its potential as a reservoir of tumor biomarkers. However, there has been less focus on the protein components of the secretory machinery itself. We therefore investigated the expression changes in secretory pathway components across many different cancer types. Specifically, we implemented a dual approach involving differential expression analysis and machine learning to identify PSP genes whose expression was associated with key tumor characteristics: mutation of p53, cancer status, and tumor stage. Eight different machine learning algorithms were included in the analysis to enable comparison between methods and to focus on signals that were robust to algorithm type. The machine learning approach was validated by identifying PSP genes known to be regulated by p53, and even outperformed the differential expression analysis approach. Among the different analysis methods and cancer types, the kinesin family members *KIF20A* and *KIF23* were consistently among the top genes associated with malignant transformation or tumor stage. However, unlike most cancer types which exhibited elevated *KIF20A* expression that remained relatively constant across tumor stages, renal carcinomas displayed a more gradual increase that continued with increasing disease severity. Collectively, our study demonstrates the complementary nature of a combined differential expression and machine learning approach for analyzing gene expression data, and highlights key PSP components relevant to features of tumor pathophysiology that may constitute potential therapeutic targets.

**Author Summary:** The secretory pathway is a series of intracellular compartments and enzymes that process and export proteins from the cell to the surrounding environment. Dysfunction of the secretory pathway is associated with many diseases, including cancer, and therefore constitutes a potential target for novel therapeutic strategies. The large number of interacting components that comprise the secretory pathway pose a challenge when attempting to identify where the dysfunction originates and/or how to restore healthy function. To improve our understanding of how the secretory pathway is changed within tumors, we used gene expression data from normal tissue and tumor samples from thousands of individuals which included many different types of cancers. The data was analyzed using various machine learning algorithms which we trained to predict sample characteristics, such as disease severity. This training quantified the relative degree to which each gene was associated with the tumor characteristic, allowing us to predict which secretory pathway components were important for processes such as tumor progression—both within specific cancer types and across many different cancer types. Our approach demonstrated excellent performance compared to traditional gene expression analysis methods and identified several secretory pathway components with strong evidence of involvement in tumor development.

## 1. Introduction

One of the most challenging features in diagnosing and treating cancer is its heterogeneity – the tissue of origin, gene mutation profile, patient, and local tumor environment are some of the many factors that can affect the pathophysiology and response to treatment of a particular cancer [1]. However, a core set of features exhibited by cancer cells establish a common thread despite other variations. Many of these shared features have been distilled into a set of “cancer hallmarks”, such as resisting cell death, activating invasion and metastasis, and avoiding immune destruction [2]. Furthermore, tumor cells acquire and sustain many of these hallmarks through interactions with each other and with neighboring “normal” cells, which together with the cancer cells form the tumor microenvironment [3]. An important system that links tumor cells to each other and to the microenvironment is the protein secretory pathway (PSP) [4]. Secreted and membrane proteins processed by the PSP contribute to critical tumor functions, such as facilitating communication among different cells residing in the microenvironment (and even with distant tissue sites in the body), and for construction and turnover of the tumor extracellular matrix. Collectively, these functions support a key role for the PSP in cancer physiology, making it an attractive target for potential therapeutic approaches.

Advancements in high-throughput molecular profiling technologies such as transcriptomics and proteomics have enabled extensive investigation and characterization of the human secretome [5] and its changes during the onset and progression of diseases such as cancer [6,7]. Although many components of the PSP that drive these important secretome changes have been studied individually, an investigation of how these constituents behave together as a system is lacking, particularly in the context of cancer. Recent efforts have begun to elucidate this system by exploring how PSP expression patterns compare to those of the secretome among different human tissues [8], and by developing genome-scale reconstructions of the PSP to mechanistically link these characteristics to the metabolic network [9]. We sought to further extend the systematic investigation of the PSP through the application of machine learning (ML) approaches.

The efficacy of ML-based approaches in the investigation of omics datasets has been demonstrated in a number of recent studies [10–14]. For example, van IJzendoorn and colleagues applied a random forest algorithm to three gene expression databases (TCGA, GTEx, and the French Sarcoma Group) to identify novel diagnostic markers for soft tissue sarcoma, which was validated with qRT-PCR in an independent experiment [11]. In another study, Wood and colleagues used L1-regularized logistic regression (Lasso) to develop a classifier for nonalcoholic fatty liver disease (NAFLD) based on phenotypic, genomic, and proteomic features [10]. Furthermore, the MLSeq R package was developed to facilitate the use of over 90 different ML algorithms for the analysis of RNA-Seq or microarray data, enabling the generation of classification models and identification of potential biomarkers [15].

We applied differential expression (DE) analysis and 8 different ML methods on RNA-seq data from The Cancer Genome Atlas (TCGA) to identify genes encoding PSP machinery that are associated with clinical features including cancer status, tumor stage, and mutation profile. The classification performance of the ML algorithms was evaluated for each of the clinical features, and relevant PSP genes were identified by DE analysis and compared with those identified by ML. The analyses reveal PSP components that exhibit pan-cancer and cancer-specific roles, and demonstrate the complementarity of DE and ML methods in the analysis of omics data.

## 2. Results

### 2.1 Data retrieval and definition of PSP genes

We retrieved 11,053 RNA-seq samples and 9,375 mutation profiles from TCGA, spanning 10,198 individuals and 33 cancer types (Table S1). Our analysis was focused on the subset of 575 genes encoding and/or regulating the human PSP machinery, as defined in the study by Feizi et al. [8]. These PSP genes encode for secretory processes such as folding, glycosylation, and trafficking, as well as protein-related stress responses (e.g., the unfolded protein response).

### 2.2 ML-based gene scoring

We developed a gene scoring approach (Fig. 1) whereby samples were grouped according to a known binary variable of interest (such as normal vs. tumor), and a ML classifier was trained to predict the group (class) of each sample based on the expression of its PSP genes. Classifiers were trained using 10-fold cross validation, and prediction performance was quantified by area under the receiver operating characteristic (ROC) curve (ROC AUC). The resulting feature importance scores of the trained classifier, which quantify roughly how useful each gene is in predicting sample class, were normalized by taking the absolute value and scaling to a range of 0 to 1. A consensus score for each gene was computed as the average score across the 8 different ML algorithms.

**Fig. 1.**
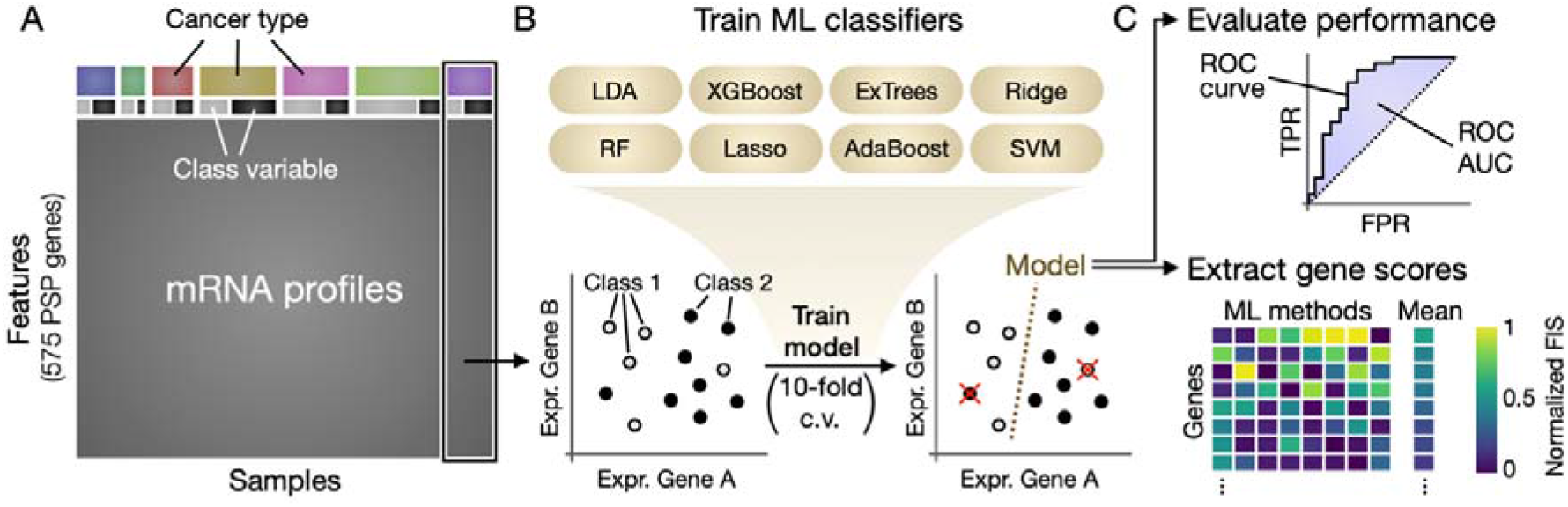
Schematic of the ML gene scoring approach. (**A**) RNA-Seq data from TCGA was filtered to remove non-PSP genes and cancer types were analyzed individually. (**B**) Samples within each cancer type were grouped according to a binary variable (e.g., Class 1 = normal; Class 2 = tumor), and 8 different ML algorithms were used to train models to predict sample class based on PSP gene expression levels (red X’s in the plot indicate failed predictions). (**C**) The prediction performance of each model was evaluated by ROC AUC, and the feature (gene) importance scores were extracted from each ML model, normalized to a range of 0-1, and averaged to obtain a consensus ML gene score. Abbreviations: c.v., cross-validation; TPR, true positive rate; FPR, false positive rate; FIS, feature importance score.

The ML algorithms used in the present study were random forests [16], extremely randomized trees (ExTrees) [17], adaptive boosting (AdaBoost) [18], extreme gradient boosted trees (XGBoost) [19], linear discriminant analysis (LDA), Lasso regression [20], Ridge regression, and support vector machine (SVM) [21]. We did not seek to include a comprehensive coverage of the available ML algorithms, as this would be infeasible and beyond the scope of the study. The algorithms were selected to include some of the most commonly employed methods for biological data [22,23], and to span different classes such as ensemble learning (random forests, ExTrees), boosting (AdaBoost, XGBoost), regularized logistic regression (lasso and ridge regression), and other common linear classifiers (LDA and SVM). Furthermore, algorithms were limited to those for which feature importance scores could be calculated.

### 2.3 Mutation of tumor protein 53

We first sought to validate our ML gene scoring approach using a class variable for which the associated gene(s) are well-established. A mutation in the *TP53* gene (encoding the p53 protein) is one of the most common mutations observed in human cancers, and the resulting loss or change in its activity as a tumor suppressor contributes to malignant progression [24]. Since p53 and its regulatory targets have been extensively characterized, we began our investigation with p53 mutation status as the class variable by which to group samples (non-mutated vs. mutated *p53*). Of the 575 PSP genes considered in the study, 3 are known to be direct targets of p53 regulation: *BCL2* Associated X (*BAX*), Heat Shock Protein Family A Member 4 Like (*HSPA4L*), and Kinesin Family Member 23 (*KIF23*). It is therefore expected that an effective approach should be able to identify some or all of these 3 genes as being associated with p53 mutation status.

Mutation data for the *TP53* gene in TCGA subjects was obtained from whole-exome sequencing data and aligned with the RNA-seq data, enabling the classification of tumor RNA-seq samples in each cancer type as “mutated” or “non-mutated” in *TP53*. Cancer types with fewer than 10 samples in each class were discarded, leaving a total of 22 different cancer types. Each of the 8 ML algorithms were trained on the data to predict p53 mutation status based on PSP gene expression, and the resulting gene scores (Table S2) and ROC AUC values (Table S3) were calculated. In addition, DE analyses were performed between mutated and non-mutated samples for each cancer type, yielding a log2 fold-change and associated significance (p-value; adjusted for the false discovery rate (FDR)) for each PSP gene (Table S4).

The consensus ML gene scores were averaged across all cancer types to identify genes that were generally associated with the p53 mutation (Fig. 2A-B). The top 3 genes were *BAX*, *HSPA4L*, and *KIF23*—precisely those known to be regulated by p53—thus providing support for the validity of the ML gene scores. Although these 3 genes were significantly differentially expressed (p53 mutated vs. non-mutated) in many of the cancer types, only *KIF23* was among the top 3 when averaging DE gene scores across all cancers, whereas *BAX* and *HSPA4L* were ranked 9th and 24th (out of 575), respectively (Fig. S1). The ML gene scoring approach thus outperformed the DE method in identifying the genes most directly associated with the p53 mutation status.

**Fig. 2.**
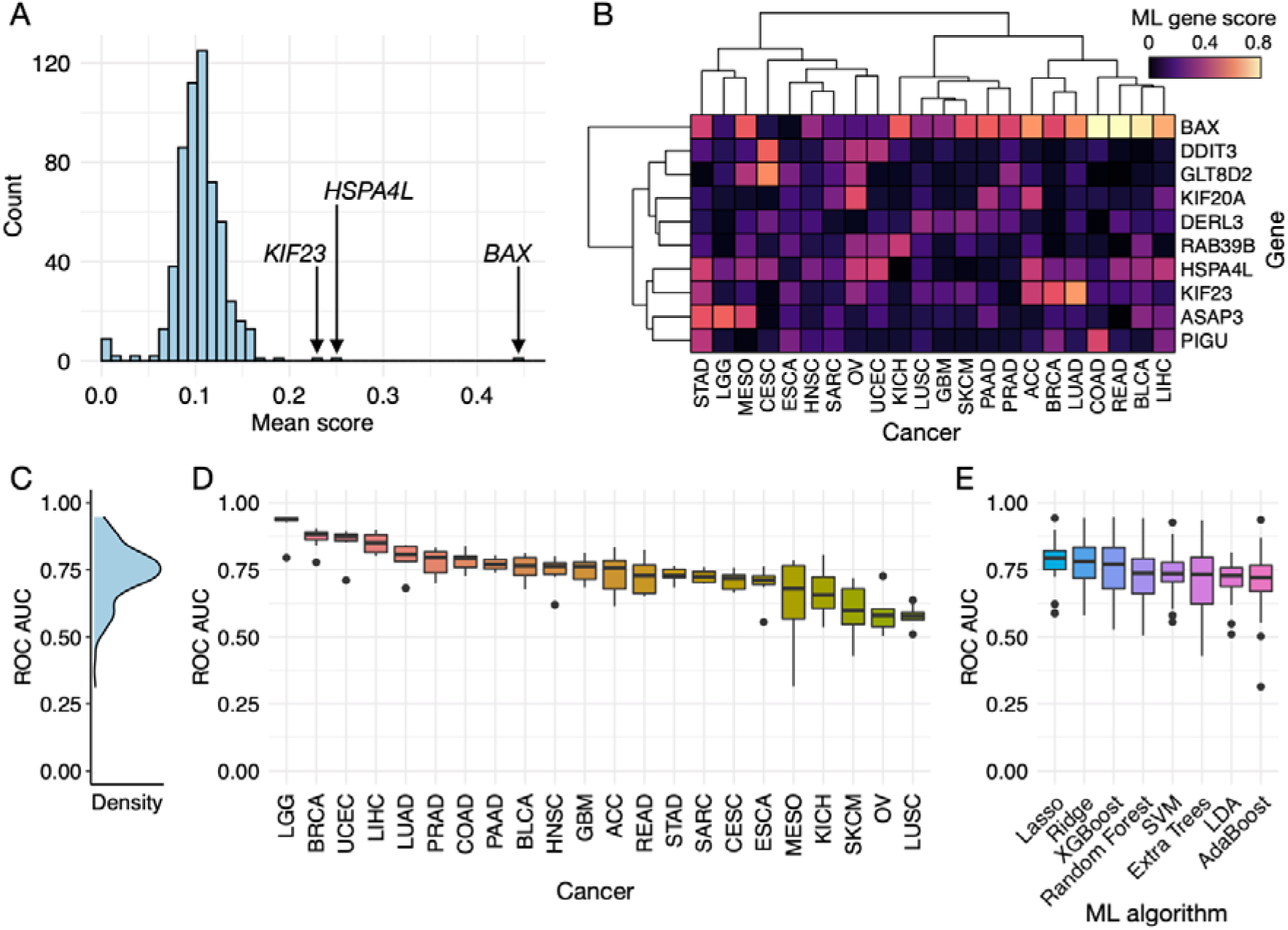
Identification of PSP genes associated with p53 mutation. (**A**) Histogram of ML gene scores averaged across all cancer types, with the top 3 scoring genes labeled. (**B**) Heatmap of the ML gene scores for each cancer type, showing only the top 10 scoring genes on average across cancer types. (**C**) Density histogram of all ROC AUC values for each ML algorithm and cancer type. Boxplots of ROC AUC values grouped by (**D**) cancer type or (**E**) ML algorithm.

Despite the excellent performance of the ML scoring approach in identifying relevant genes, this was not entirely reflected in the associated ROC AUC values. The average ROC AUC across all cancer and ML algorithm types was 0.74 ± 0.11 (mean ± standard deviation) (Fig 2C), where a value of 1 corresponds to a perfect predictor and 0.5 is no better than random. There were no clear differences between algorithms in terms of ROC AUC, though the regularized regression methods (Ridge and Lasso) exhibited slightly higher values. A much larger difference was observed between cancer types, where the average ROC AUC for LGG, BRCA, and UCEC exceeded 0.85, but was at or below 0.60 for SKCM, OV, and LUSC (Fig. 2D-E). Interestingly, *BAX*, *HSPA4L*, and *KIF23* were not among the top-scoring genes for any of the three cancer types with the highest ROC AUC values, except for *KIF23* in BRCA. This suggests that useful information can be extracted from the feature scores of trained ML classifiers despite a relatively poor corresponding ROC AUC.

### 2.4 Investigation of PSP genes associated with malignant transformation

After validating the ML gene scoring approach, we used it to evaluate the relative importance of each PSP component in distinguishing normal vs. tumor samples and identify genes that are likely to contribute to the tumor phenotype. Analogous to the p53 mutation analysis, samples for each cancer type were grouped according to cancer status (normal or tumor) and each of the 8 ML algorithms, as well as DE analysis, were used to score the 575 PSP genes. Cancer types without at least 10 samples in each group were excluded, yielding a total of 16 cancer types.

Unlike the classifiers trained on p53 mutation status, the ROC AUC for predicting normal vs. tumor samples based on PSP gene expression was high across all cancer types and ML methods, with an overall average of 0.98 ± 0.03 (Fig. S2). Only the LDA algorithm and the ESCA cancer type tended to exhibit lower ROC AUC values relative to the others, but the lowest value for each was still greater than 0.80. This higher prediction performance for cancer status as compared to p53 mutation status was expected since there is a much broader range of differences between normal and tumor tissues than there are between tumor cells differing in a single gene mutation.

#### 2.4.1 Pan-cancer features

Inspection of the ML gene scores and DE analysis results revealed that kinesin-6 family proteins (*KIF20A* and *KIF23*), Crystallin Alpha B (*CRYAB*), and a few proteins belonging to the soluble N-ethylmaleimide-sensitive-factor attachment protein receptor (*SNARE*) family (*STX1A*, *STX12*, *STX11*, and *VAMP2*) generally scored highly in both ML and DE approaches among the different cancer types (Fig. 3A-B, Fig. S3), suggesting that these proteins play an important role in tumor physiology. *KIF20A* and *KIF23* were among the top 3 genes with the highest average ML consensus scores and exhibited a significant (FDR adjusted p-value < 0.01) expression increase in tumor compared to normal samples for all 16 cancer types except two renal carcinomas, KICH and KIRP (Fig. S4). Although they are associated with Golgi-to-ER retrograde transport and intracellular organelle transport, *KIF20A* and *KIF23* play a critical role in mitosis and cytokinesis [25,26]. Inhibitors of these and other kinesin family proteins are undergoing clinical trials as anticancer therapeutics [27].

**Fig. 3.**
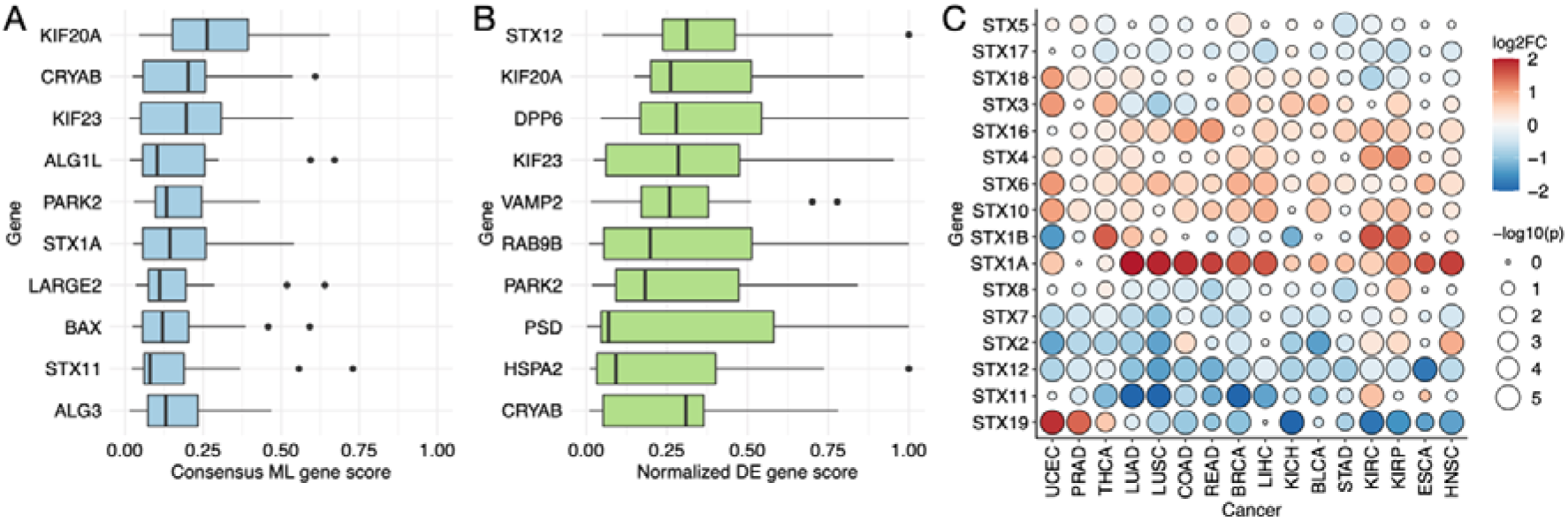
Kinesins and components of the SNARE complex are associated with cancer status. Boxplots show the (**A**) consensus ML gene scores and (**B**) normalized DE gene scores among the different cancer types. Only the top 10 scoring genes on average for each scoring type are shown. (**C**) Log-transformed expression fold-changes and significance (FDR-adjusted p-values) of PSP genes belonging to the STX family, from the DE analysis. Color indicates fold-change magnitude and direction, whereas circle size indicates significance.

Although *STX1A* exhibited a similar expression increase across most cancer types as the kinesin-6 family proteins, the expression of *STX11*, *STX12*, and *VAMP2* was significantly decreased in tumors across nearly all 16 cancer types. We further investigated the expression changes of the PSP genes belonging to the *STX* (Fig. 3C) or *VAMP* (Fig. S5) gene families. There was a common restructuring pattern of *STX* expression among the different cancer types, involving a mixture of increases and decreases across the different *STX* genes, whereas *VAMP* genes tended to be more broadly decreased with the exception of *VAMP1* and *VAMP8*. *SNARE* proteins, which include the *STX* and *VAMP* families, mediate the membrane fusion necessary for trafficking through the different steps of the secretory pathway [28]. *SNARE*s have been found to support many tumorigenic functions such as autophagy, cell invasion, and chemo-resistance, and thus constitute potential targets in anti-cancer therapies [29].

The *CRYAB* gene exhibited the second highest ML consensus score on average across the 16 cancer types (Fig. 3A), and was significantly differentially expressed (FDR-adjusted p < 0.01) in all but 3 cancer types. Unlike the kinesins whose expression was nearly always increased in tumor relative to normal tissue, *CRYAB* expression was significantly decreased in tumor for 10 cancer types and increased in only 3: LIHC, KIRC, and KIRP (Fig. S6). The mean *CRYAB* mRNA abundance of LIHC samples were the lowest of all cancer types (< 10 TPM) and thus the DE results are less reliable; however, both KIRC and KIRP exhibited among the highest expression of *CRYAB* in paired normal samples which further increased by 1.9- and 6.5-fold in their corresponding tumor samples, respectively. The main role of *CRYAB* is to form multimeric structures with other proteins to prevent aggregation, but it has also been shown to exhibit other activities such as protection from oxidative stress and apoptotic stimuli [30]. In the context of cancer, there does not appear to be a clear consensus as to whether *CRYAB* supports or suppresses tumorigenesis [30]. Many studies conclude a pro-tumorigenic effect of *CRYAB* and a positive correlation between its expression and tumor aggression [31], whereas others report a tumor-suppressive activity and/or decreased expression in more aggressive tumors [32,33]. Our results suggest that cancer type is one factor determining whether *CRYAB* exerts an inhibitory or supportive role in a tumor, and that renal carcinomas in particular may be susceptible to *CRYAB*-modulating therapies.

#### 2.4.2 Cancer-specific features

Although the kinesins, *SNARE*s, and *CRYAB* were among the highest ML gene scores when averaging over all 16 cancer types, no genes were consistently high-scoring in more than a few of the cancer types. An inspection of the top-scoring genes of each individual cancer type revealed that high-scoring genes were primarily cancer-specific (Fig. S7). For example, *RAS* oncogene family member 17 (*RAB17*) scored highly in prostate adenocarcinoma (PRAD) across nearly all ML algorithms with a consensus score of 0.82, whereas its score in all other cancer types ranged from 0.01 to 0.17. Members of the *RAB* family regulate vesicle trafficking and are known to both promote and suppress tumor growth, depending on the family member and cancer type [34]. Although increased expression of *RAB25* has been shown to contribute to prostate cancer malignancy and recurrence [35], similar studies or observations involving *RAB17* are lacking.

When investigating the highest-scoring genes for each individual cancer type, we observed a high frequency of genes associated with glycosylation, particularly for five cancer types: STAD, READ, COAD, KICH, and THCA (Fig 4). For each of these cancer types, 3 out of their top 5 scoring genes encoded some form of glycosylation activity, despite such activity accounting for less than 18% of the 575 PSP genes considered in this study.

**Fig. 4.**
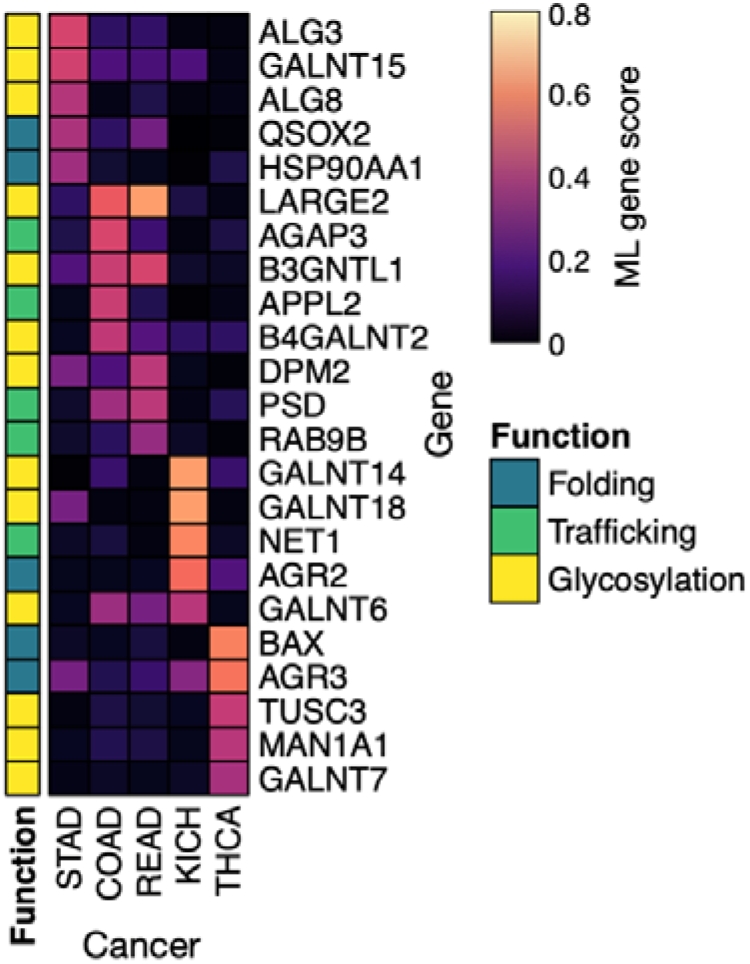
Glycosylation is an enriched function among the top PSP genes associated with a subset of cancer types. The heatmap shows the consensus ML gene scores of the cancer types for which 3 out of 5 top-scoring genes encode for glycosylation activity. The colorbar on the left indicates the function associated with each gene.

Genes associated with O-linked peptide glycosylation (*LARGE2*, *B3GNTL1*, *B4GALNT2*, and the *GALNT* family) were associated with KICH, COAD, and to a lesser extent READ, whereas genes encoding N-linked glycosylation activity (*DPM2*, *MAN1A1*, *TUSC3*, and the *ALG* family) scored highly for the STAD and THCA cancer types. The expression and specific patterns of glycans dictate cellular functions such as adhesion, signal transduction, differentiation, and proliferation, and the alteration of such patterns is a hallmark of tumor physiology [36,37]. It is therefore logical that genes encoding these post-translational modifications scored highly in the ML classifiers distinguishing normal from tumor samples. Furthermore, the cancer-specificity of these high-scoring genes is likely a reflection of the specificity and complexity of the glycosylation machinery and its large repertoire of glycan patterns [38].

### 2.5 Analysis of different tumor stages

We next focused on PSP gene expression changes between tumor stages to identify secretory pathway components that were associated with disease severity and tumor development. Most TCGA samples are annotated with tumor stage information which generally ranges from stage I to stage IV, enabling the investigation of transcriptomic changes as a function of disease progression. Primary tumor samples were grouped into stages I, II, III, and IV within each cancer type, and ML classifiers were trained on all possible pairs of tumor stages to predict a sample’s stage based on its corresponding PSP gene expression profile. Cancer types without at least 10 samples in at least 3 tumor stages were discarded, yielding a total of 17 cancer types.

The ROC AUC values for the ML classifiers of tumor stage were substantially lower than those trained to separate normal vs. tumor samples, where many performed no better than random (ROC AUC ~ 0.5) (Fig. 5A and Fig. S8). This was expected given that physiological differences between tumor stages are relatively subtle when compared to those between normal and cancerous tissue. Although our initial analysis with the p53 mutation ML classifiers suggested that feature importance scores can still provide some meaningful information despite relatively low ROC AUC values, we expect that such scores will largely degrade into random noise when approaching very poor values near (or below) 0.5. We therefore continued the analysis using only the top 5 performing cancer types based on their average ROC AUC values across the different ML algorithms and pairs of tumor stages: THCA, TGCT, KIRP, KIRC, and ACC (Fig. 5A).

**Fig. 5.**
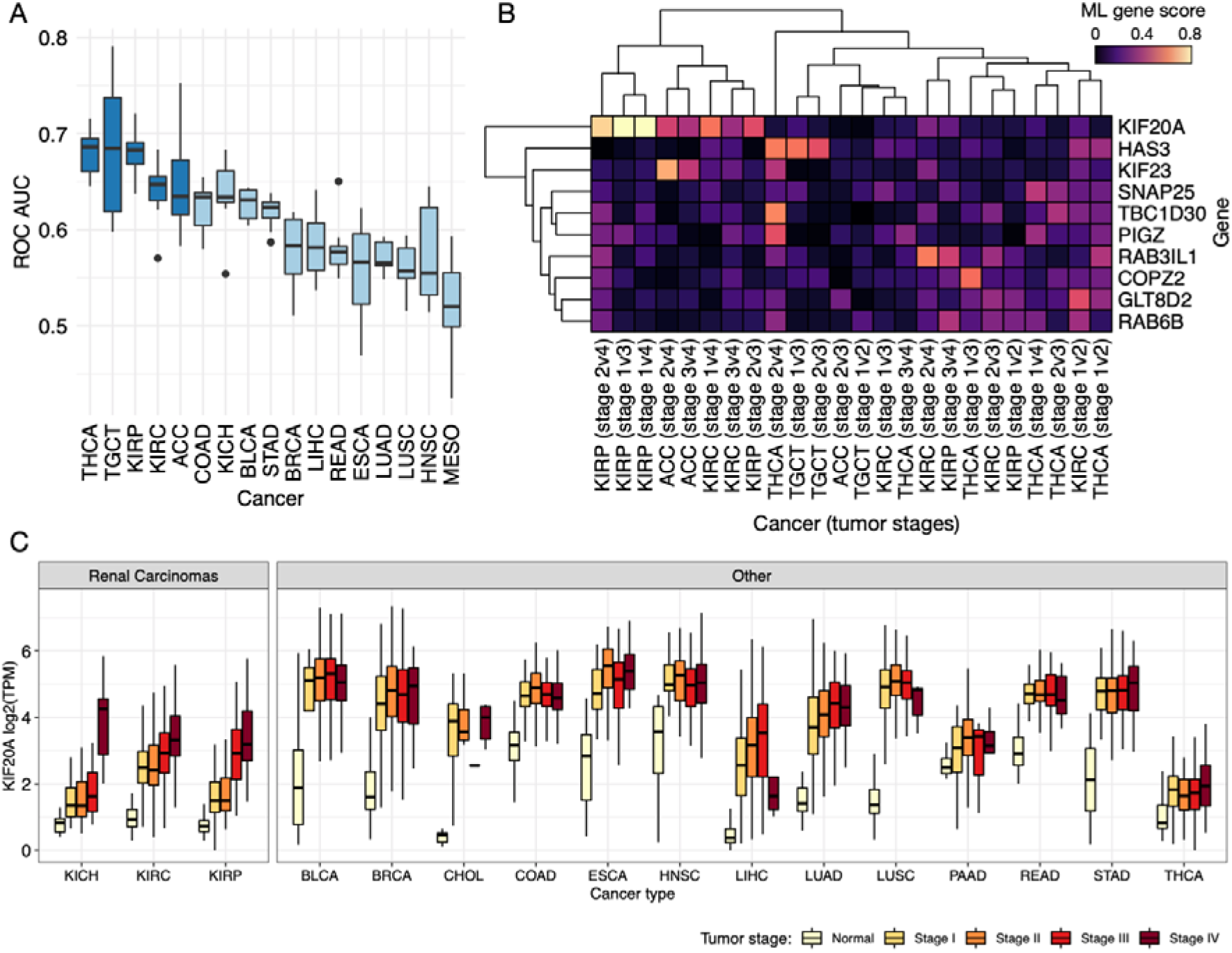
PSP genes associated with tumor stage. (**A**) Boxplots of mean ROC AUC values for the prediction of tumor stage based on PSP gene expression. Only cancer types with sufficient samples in >2 different tumor stages are included. Dark blue boxes indicate the cancer types with the top 5 ROC AUC values on average, which were used in subsequent analyses. (**B**) Heatmap of consensus ML gene scores for the different stage comparisons of each cancer type, showing only the top 10 scoring genes on average. (**C**) Expression (log transformed TPM) of *KIF20A* among different cancer types, grouped by tumor stage.

Among the PSP genes exhibiting the highest average consensus ML scores across the different cancer types and tumor stages, hyaluronan synthase 3 (*HAS3*) and *KIF20A* were the most prominent (Fig. 5B and Fig. S9). Hyaluronan, an extracellular matrix polysaccharide, is enriched in the matrix surrounding virtually all epithelial tumors [39], and has been shown to promote tumor malignancy and metastasis by increasing cell invasiveness and anchorage-independent growth [40]. Consistent with the malignant function of hyaluronan, *HAS3* exhibited a high consensus ML score for TGCT and to a lesser extent THCA, suggesting an association between *HAS3* expression and tumor stage. This was reflected in the gene expression profiles of TGCT tumor samples, which showed increasing expression of *HAS3* and *HAS2* with increasing tumor stage (Fig. S10). Conversely, THCA did not exhibit a substantial difference in *HAS3* expression as a function of tumor stage, but a positive relationship was observed between THCA tumor stage and *HAS1* expression (Fig. S10).

The top-scoring gene on average across the 5 cancer types was *KIF20A*, which exhibited particularly high scores for distinguishing tumor stages of KIRP, and to a lesser extent those of KIRC and ACC. Interestingly, the renal carcinomas (KIRP, KICH, and KIRC) were among the few cancer types for which the kinesins (*KIF23* and *KIF20A*) were either not significantly DE and/or exhibited a very low consensus ML score (less than 0.1 on average) when comparing normal to tumor tissues. This distinction becomes clear when comparing the expression of *KIF20A* between normal tissue and different tumor stages for each cancer type (Fig. 5C). Most cancer types exhibit a sharp increase in *KIF20A* expression between normal and tumor samples that remain relatively constant across the different tumor stages, whereas renal carcinomas display a more gradual change in *KIF20A* expression that continues to increase with increasing tumor stage. Although *KIF20A* has been implicated in the development of many other cancer types [27,41,42], its involvement in renal carcinoma has not been addressed. The expression dynamics observed here suggest that *KIF20A* may support more invasive and metastatic functions associated with later stages of renal carcinoma, and thus constitutes a potential therapeutic target for this cancer type.

## 3. Discussion

The secretory pathway and its products are essential to the viability of eukaryotic organisms, but the deregulation of secretory machinery can support detrimental processes such as those driving tumorigenesis [7,43]. The identification of PSP components exhibiting oncogenic or tumor suppressive activities can aid in the development of novel anti-cancer therapies that aim to restore healthy PSP function through the modulation of these components. We therefore conducted a focused investigation of the PSP transcriptional changes associated with malignant transformation and tumor progression across many different cancer types. This allowed us to identify patterns in PSP expression that were common to carcinogenesis independent of cancer type, as well as explore secretory elements that exhibited cancer type specific behavior.

Accessing and interpreting the information embedded within omics data is non-trivial due to its high volume and dimensionality, and has traditionally been limited to a few methods, such as DE analysis and principal component analysis (PCA) [44]. We therefore sought to deepen the investigation by applying different machine learning (ML) approaches to provide a more detailed understanding of PSP behavior in tumors. However, ML methods generally struggle when the number of features (genes) greatly exceeds the number of samples, which is often the case for RNA-seq or other omics datasets and is referred to as the curse of dimensionality [45]. The ML methods were therefore well-suited for this focused study because the number of features was greatly reduced by including only the 575 PSP genes in the analyses. Furthermore, there is often a risk of over-fitting or a high frequency of false positives when using a data-driven approach such as ML. We therefore implemented 8 different types of ML algorithms in our analyses and used a normalized average consensus score that combined the results of each algorithm. This ensured that genes identified as relevant to a given biological class variable were robust to the choice of ML algorithm or effects of overfitting.

The use of several different ML algorithms also enabled comparison of their predictive performance for each of the investigations, as quantified by ROC AUC. Although some ML algorithms (such as regularized regression and extreme gradient boosted trees) tended to outperform others (such as LDA and adaptive boosting) among the different class variables and cancer types, the difference was marginal and far from significant. This further supported using a consensus score that combined the output of the 8 different methods with equal weighting because no method consistently outperformed the others.

We used a well-studied feature in cancer biology - the mutation of p53 - to evaluate the performance of the ML approach in terms of identifying biologically relevant features, and to compare with DE analysis. The highest consensus ML gene scores were exhibited by the known regulatory targets of p53 in the 575 PSP genes (*BAX*, *HSPA4L*, and *KIF23*), providing confidence in the biological relevance of the ML results. Although the DE analysis identified these genes as important, some were not ranked as highly as other PSP genes. A reason for why the ML methods outperformed DE analysis in this case is because the ML algorithms can capture interactions between genes and their expression patterns in different samples, whereas DE analysis estimates a fold-change and confidence for each gene individually.

A recurring gene of importance in our analyses was *KIF20A*, which was remarkably among the top-scoring genes for both ML and DE approaches and for all class variables (p53 mutation status, normal vs. tumor, and different tumor stages). This is consistent with the abundance of studies that have identified *KIF20A* to be highly expressed, linked to tumor aggressiveness, correlate with poor survival, diagnostic, and/or prognostic in many different cancer types [41], which support a critical and diverse role of the protein in general tumor development and progression. There are however a lack of studies identifying any role or association of *KIF20A* with renal carcinoma other than a co-expression network analysis of clear cell renal cell carcinoma (ccRCC) by Yuan and colleagues, in which *KIF20A* was identified as one of six hub genes associated with ccRCC progression [46]. We observed a substantial difference in the pattern of *KIF20A* expression among normal and stage-stratified tumor samples in all renal carcinomas (KICH, KIRP, KIRC) compared to other cancer types; *KIF20A* expression in renal carcinomas increased more gradually with increasing tumor stage rather than a sharp increase between normal and tumor that remained relatively constant across stages. We cannot speculate from this data alone as to the cause for the different dynamics, but it may indicate that anti-cancer treatments targeting *KIF20A* could exhibit variable efficacy with tumor stage for renal carcinomas.

Our investigation demonstrates the efficacy of using a consensus ML-based gene scoring approach to predict biologically relevant features from a focused set of genes, and highlights the utility of using such an approach to complement and support the results of DE analysis. Furthermore, we present a set of PSP-associated proteins and protein families that exhibit a robust association with malignant transformation and tumor progression, and thus hold potential as targets in the development of anti-cancer therapeutics.

## 4. Methods

### Analysis and figure scripts

The scripts used to perform the analyses and generate the figures presented here, as well as all analysis outputs, are available on GitHub: https://github.com/SysBioChalmers/CancerProteinSecretionML. Data files too large to host on GitHub were deposited on Zenodo: https://doi.org/10.5281/zenodo.3978373.

### RNA-Seq and mutation data retrieval

Transcriptomic (RNA-seq) and mutation annotation data was retrieved from TCGA using the TCGAbiolinks R package [47]. Raw gene counts and normalized (FPKM) gene counts were retrieved for 33 available cancer types. Mutation annotation information was obtained using the MuTect2 variant calling pipeline [48], and processed such that each gene in each sample was classified as mutated if it was modified in any way (insertion, deletion, missense, silent, etc.), otherwise it was classified as non-mutated.

### Differential expression analysis

Differential expression analysis was performed on raw gene counts using the edgeR package [49]. Samples were grouped according to a binary class variable of interest (e.g., p53 mutation status, cancer status, or tumor stage), and all genes were included except for those that had fewer than 10 counts in 70% of the samples of the smallest group. Genes excluded from an analysis due to low counts were automatically assigned a log2 fold-change of zero and a p-value of one. The design matrix included only information regarding group membership of each sample. We did not account for patient identity when performing the normal vs. primary tumor analysis because it would require the exclusion of many tumor samples which were used in the ML analyses. The expression fold-changes and associated significance (FDR-adjusted p-values) were calculated prior to filtering out non-PSP genes. Analyses were performed on each cancer type individually, where cancer types with fewer than 10 samples in a group were excluded.

A DE gene score was formulated to enable comparison with the ML gene score described below. The FDR-adjusted p-values of PSP genes for a given comparison and cancer type were log-transformed, negated, and normalized to a range of 0 to 1.

### ML model training and gene scoring

All ML methods were implemented in python using the scikit-learn package [50] or the XGBoost package [19]. Gene expression values were converted to transcripts per million (TPM), and natural log transformed after adding a pseudocount of 1 TPM to avoid logarithm of zero. Samples were grouped according to a binary class variable of interest (e.g., p53 mutation status, cancer status, or tumor stage), and cancer types were analyzed individually, where cancer types with fewer than 10 samples in a group were excluded. Non-PSP genes and genes with a median expression below 0.1 TPM among both sample groups were also excluded.

For each cancer type and class variable, 8 classification models were trained using each of the 8 ML algorithms (random forests, ExTrees, AdaBoost, XGBoost, LDA, lasso regression, ridge regression, and SVM). Default parameters were used for each algorithm when available. For the tree-based methods, the number of estimators was set to the recommended value of the square root of the number of features, rounded down to the nearest integer. For the ridge and lasso regression methods, the “saga” solver was used with a maximum of 10,000 iterations. Training was performed using stratified 10-fold cross validation, such that each fold contained approximately the same proportion of samples from each group. Feature importance scores were extracted from each trained model, and normalized by taking the absolute value and scaling to a range of 0 to 1. A consensus score was calculated for each gene by averaging the normalized importance scores obtained from each of the 8 algorithms.

We note that our primary interest was to determine the relative importance of features (genes), as quantified by the gene scores, rather than developing predictive models. We therefore used all available samples when training each classifier and did not exclude any samples for a separate test set, meaning that the reported ROC AUC values are likely higher than what one would expect if the trained model predictions were evaluated using an independent test set of samples. The ROC AUC values were determined using stratified 10-fold cross validation, where the reported values are the mean of the 10 folds.

### Tumor stage processing

Tumor stages in TCGA are often provided with sub-stage detail, such as stage IIa, stage IIb, etc. We merged such annotations to achieve only four different stages: I, II, III, and IV. The merging was performed to avoid increasingly large numbers of pairwise stage comparisons, as well as groups with very few samples.

## Acknowledgements

Research reported in this publication was supported by funding from the Knut and Alice Wallenberg Foundation, and from the Information and Communication Technology and Life Science Areas of Advance at Chalmers University of Technology. JLR is financially supported by the Knut and Alice Wallenberg Foundation as part of the National Bioinformatics Infrastructure Sweden at SciLifeLab.

## Author Contributions

Conceptualization: JN and JLR; methodology: RS, ASM, PB, and JLR; investigation: RS, ASM, and JLR; writing (original draft): RS and JLR; writing (review and editing): ASM, PB, and JN; supervision: JN and JLR; funding acquisition: JN and JLR.

## Supporting information captions

### Supporting Figures

**Figure S1. Consensus ML and DE gene scores for p53 mutation status.** Histogram of (**A**) mean ML gene scores and (**B**) mean DE gene scores across all available cancer types, where the three PSP genes known to be regulated by p53 are labeled. Boxplots of (**C**) consensus ML gene scores and (**D**) DE gene scores for the top 10 scoring genes on average. Clustered heatmaps showing the (**E**) consensus ML gene scores and (**F**) DE gene scores for individual cancers for the top 10 scoring genes on average.

**Figure S2. ROC AUC values for the prediction of cancer status by the trained ML models.** (**A**) Density histogram of all ROC AUC values across different cancer types and ML algorithms. Boxplots showing the ROC AUC values grouped by (**B**) cancer type or (**C**) ML algorithm.

**Figure S3. Consensus ML and DE gene scores for cancer status.** Histogram of (**A**) mean ML gene scores and (**B**) mean DE gene scores across all available cancer types. Boxplots of (**C**) consensus ML gene scores and (**D**) DE gene scores for the top 10 scoring genes on average. Clustered heatmaps showing the (**E**) consensus ML gene scores and (**F**) DE gene scores for individual cancers for the top 10 scoring genes on average.

**Figure S4. Expression fold-change and significance of PSP genes belonging to the KIF family from the DE analysis of normal vs. tumor.** Color indicates fold-change magnitude and direction, whereas circle size indicates significance (FDR-adjusted p-value).

**Figure S5. Expression fold-change and significance of PSP genes belonging to the VAMP family from the DE analysis of normal vs. tumor.** Color indicates fold-change magnitude and direction, whereas circle size indicates significance (FDR-adjusted p-value).

**Figure S6. Expression of CRYAB in normal and tumor tissue samples across different cancer types.** Cancer types are grouped according to whether CRYAB significantly (FDR-adjusted p-value < 0.01) changed in expression between normal and tumor, and whether that change was a decrease or increase.

**Figure S7. Heatmap of the consensus ML gene scores for cancer status.** The heatmap includes all available cancer types and the top 5 scoring genes of each type. For visual aid, rows and columns were clustered such that high-scoring genes for each cancer tend to lie along or near the diagonal.

**Figure S8. ROC AUC values for the prediction of tumor stage by the trained ML models.** (**A**) Density histogram of all ROC AUC values across different cancer types and ML algorithms. Boxplots showing the ROC AUC values grouped by (**B**) cancer type, (**C**) ML algorithm, or (**D**) each pair of tumor stages.

**Figure S9. Consensus ML and DE gene scores for tumor stage.** Histogram of (**A**) mean ML gene scores and (**B**) mean DE gene scores across the 5 cancer types with the highest average ROC AUC values. Boxplots of (**C**) consensus ML gene scores and (**D**) DE gene scores for the top 10 scoring genes on average. Clustered heatmaps showing the (**E**) consensus ML gene scores and (**F**) DE gene scores for individual cancers for the top 10 scoring genes on average.

**Figure S10. Expression of PSP genes belonging to the HAS family among different tumor stages in TGCT and THCA cancer types.**

### Supporting Tables

**Table S1. TCGA cancer abbreviations and sample metadata.** [.docx]

**Table S2. Consensus ML gene scores for p53 mutation, cancer status, and tumor stage.** [.xlsx]

**Table S3. ROC AUC values of each ML algorithm for predicting p53 mutation, cancer status, and tumor stage.** [.xlsx]

**Table S4. Differential expression log2 fold-changes, FDR-adjusted p-values, and gene scores for p53 mutation, cancer status, and tumor stage.** [.xlsx]

