## Supplemental Table 1 for "Machine learning-based investigation of the cancer protein secretory pathway"

Table S1. TCGA cancer abbreviations and sample metadata.

|  |  | **Cancer Status** | | **TP53 Mutated** | | **Tumor Stage** | | | |
| --- | --- | --- | --- | --- | --- | --- | --- | --- | --- |
| **Code** | **Name** | Solid Tissue Normal | Primary solid Tumor | False | True | Stage I | Stage II | Stage III | Stage IV |
| ACC | Adrenocortical Carcinoma | 0 | 79 | 65 | 14 | 9 | 37 | 16 | 15 |
| BLCA | Bladder Urothelial Carcinoma | 19 | 414 | 216 | 195 | 4 | 134 | 149 | 144 |
| BRCA | Breast Invasive Carcinoma | 113 | 1101 | 641 | 338 | 202 | 697 | 276 | 22 |
| CESC | Cervical Squamous Cell Carcinoma and Endocervical Adenocarcinoma | 3 | 304 | 264 | 22 | 0 | 0 | 0 | 0 |
| CHOL | Cholangiocarcinoma | 9 | 36 | 33 | 3 | 26 | 10 | 1 | 8 |
| COAD | Colon Adenocarcinoma | 41 | 476 | 184 | 219 | 85 | 209 | 140 | 73 |
| DLBC | Lymphoid Neoplasm Diffuse Large B-cell Lymphoma | 0 | 48 | 32 | 5 | 0 | 0 | 0 | 0 |
| ESCA | Esophageal Carcinoma | 11 | 161 | 29 | 131 | 21 | 71 | 51 | 8 |
| GBM | Glioblastoma Multiforme | 0 | 155 | 100 | 48 | 0 | 0 | 0 | 0 |
| HNSC | Head and Neck Squamous Cell Carcinoma | 44 | 500 | 154 | 340 | 27 | 86 | 86 | 278 |
| KICH | Kidney Chromophobe | 24 | 65 | 45 | 20 | 29 | 33 | 17 | 10 |
| KIRC | Kidney Renal Clear Cell Carcinoma | 72 | 538 | 325 | 9 | 297 | 70 | 139 | 102 |
| KIRP | Kidney Renal Papillary Cell Carcinoma | 32 | 288 | 273 | 5 | 187 | 23 | 63 | 19 |
| LAML | Acute Myeloid Leukemia | 0 | 0 | 0 | 0 | 0 | 0 | 0 | 0 |
| LGG | Brain Lower Grade Glioma | 0 | 510 | 273 | 229 | 0 | 0 | 0 | 0 |
| LIHC | Liver Hepatocellular Carcinoma | 50 | 371 | 254 | 105 | 191 | 98 | 97 | 6 |
| LUAD | Lung Adenocarcinoma | 59 | 533 | 264 | 254 | 324 | 136 | 97 | 28 |
| LUSC | Lung Squamous Cell Carcinoma | 49 | 502 | 89 | 401 | 271 | 179 | 89 | 8 |
| MESO | Mesothelioma | 0 | 86 | 68 | 13 | 10 | 16 | 44 | 16 |
| OV | Ovarian Serous Cystadenocarcinoma | 0 | 374 | 25 | 247 | 0 | 0 | 0 | 0 |
| PAAD | Pancreatic Adenocarcinoma | 4 | 177 | 63 | 107 | 21 | 150 | 3 | 5 |
| PCPG | Pheochromocytoma and Paraganglioma | 3 | 178 | 178 | 0 | 0 | 0 | 0 | 0 |
| PRAD | Prostate Adenocarcinoma | 52 | 498 | 436 | 56 | 0 | 0 | 0 | 0 |
| READ | Rectum Adenocarcinoma | 10 | 165 | 30 | 102 | 34 | 53 | 53 | 26 |
| SARC | Sarcoma | 2 | 259 | 145 | 90 | 0 | 0 | 0 | 0 |
| SKCM | Skin Cutaneous Melanoma | 1 | 103 | 398 | 67 | 77 | 140 | 171 | 24 |
| STAD | Stomach Adenocarcinoma | 32 | 375 | 197 | 175 | 59 | 126 | 156 | 42 |
| TGCT | Testicular Germ Cell Tumors | 0 | 134 | 127 | 1 | 56 | 12 | 14 | 0 |
| THCA | Thyroid Carcinoma | 58 | 502 | 485 | 2 | 321 | 59 | 125 | 61 |
| THYM | Thymoma | 2 | 119 | 114 | 4 | 0 | 0 | 0 | 0 |
| UCEC | Uterine Corpus Endometrial Carcinoma | 23 | 551 | 331 | 199 | 0 | 0 | 0 | 0 |
| UCS | Uterine Carcinosarcoma | 0 | 56 | 4 | 52 | 0 | 0 | 0 | 0 |
| UVM | Uveal Melanoma | 0 | 80 | 80 | 0 | 0 | 39 | 36 | 4 |
